# Tree inference for single-cell data

**DOI:** 10.1101/047795

**Authors:** Katharina Jahn, Jack Kuipers, Niko Beerenwinkel

**Author notes:** Equal contributors (Katharina Jahn, Jack Kuipers).

## Abstract

Understanding the mutational heterogeneity within tumours is a keystone for the development of efficient cancer therapies. Here, we present SCITE, a stochastic search algorithm to identify the evolutionary history of a tumour from noisy and incomplete mutation profiles of single cells. SCITE comprises a exible MCMC sampling scheme that allows the user to compute the maximum-likelihood mutation history, to sample from the posterior probability distribution, and to estimate the error rates of the underlying sequencing experiments. Evaluation on real cancer data and on simulation studies shows the scalability of SCITE to present-day single-cell sequencing data and improved reconstruction accuracy compared to existing approaches.

## Background

Tumour progression can be described as a dynamic evolutionary process acting at the level of individual cells (Merlo et al., 2006; Nowell, 1976; Pepper et al., 2009). A tumour typically arises from a single founder cell whose distinct set of genetic (and epigenetic) lesions gives it a growth advantage over the surrounding cells and helps it to evade the patient’s immune response. As a consequence, the clone arising from this cell expands and, over the course of time, the descendant cells develop further into subclones by acquiring additional somatic mutations (Nik-Zainal et al., 2012). The subclones compete against each other for resources in the tumour environment and the more successful ones will replace others until eventually they themselves are out-competed by new subclones (Nik-Zainal et al., 2012; Yates and Campbell, 2012); see also Fig. 1(a).

**Figure 1.**
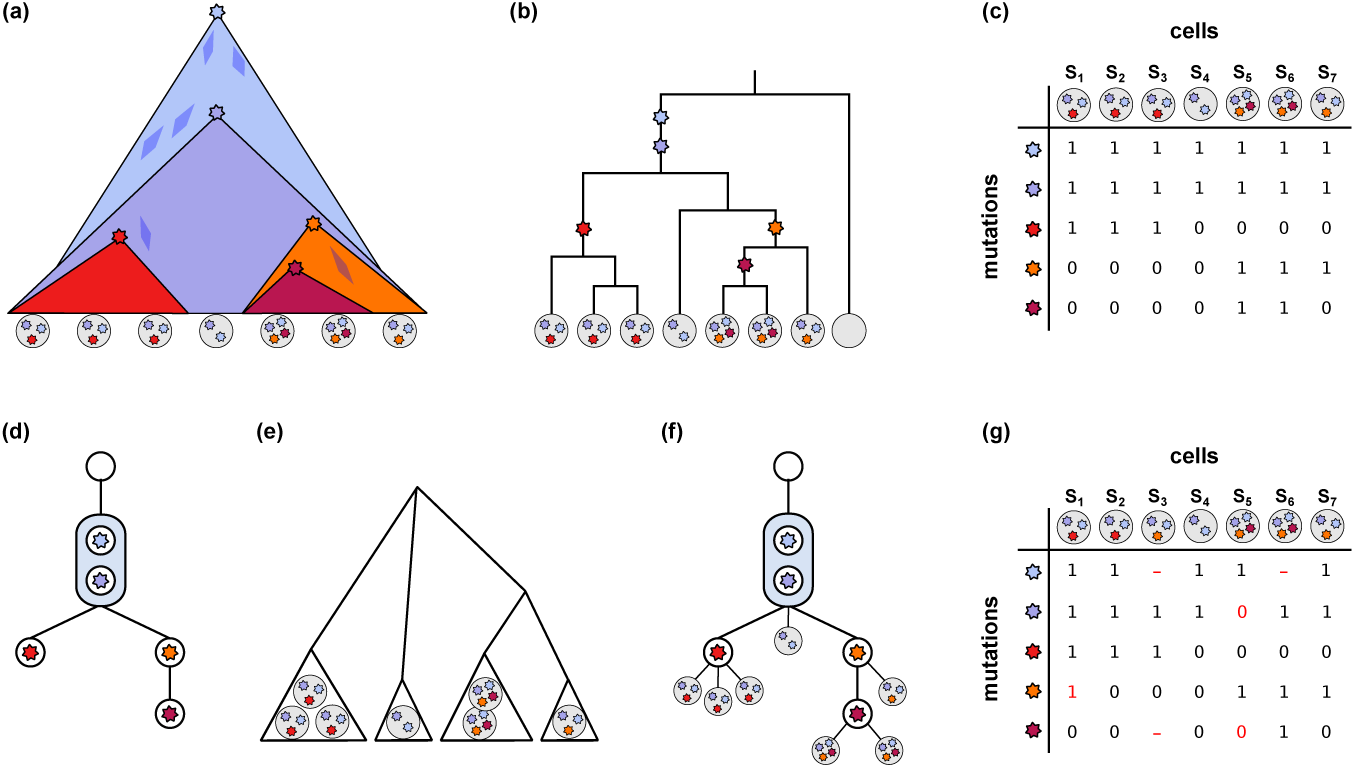
Tumour evolution and cell phylogeny. (a) Schematic representation of tumour evolution with time progressing downwards. Stars denote new mutations leading to subclone expansion. The quadrangles belong to minor extinct subclones with no traces in the present day populations. The mutations founding these clones may not have induced a sufficient growth advantage to have surviving descendant cells or may have been lost by chance. The grey discs on the bottom denote single cells sequenced after tumour removal. The stars they contain indicate the mutations observed in the cell. (b) Binary genealogical tree of the sequenced cells. Empty disc represents a normal somatic cell which is an outgroup for the tumour cells. (c) Binary mutation matrix representing the mutation status of the sequenced tumour cells. A zero entry denotes the absence of a mutation in the respective cell, a one denotes its presence. (d) The perfect phylogeny represented as a mutation tree, the partial (temporal) order of the mutation events. Mutations are summarized in a single node when their order is unidentifiable from the sampled cells, as is the case here for the two top-most mutations with the matrix from subfigure (c). (e) Hierarchical subclone structure: Cells with identical mutation profiles cluster into subclones which serve as taxa in this phylogenetic tree. (f) Mutation tree with single cell samples attached. (g) Noisy mutation matrix with missing values. The red numbers indicate flipped mutation states with respect to the true mutation matrix in (c): 0 → 1, a false positive, the mutation is called but not present in the cell; 1 → 0, a false negative, the mutation is not called but present in the cell, most likely due to allelic dropout during the DNA amplification. The red dash indicates a missing value; it is unknown whether the site is mutated or in the normal state in this cell.

The genetic diversity arising from this process, referred to as intra-tumour heterogeneity, is believed to be a major cause of relapse after cancer treatment (Ding et al., 2012; Greaves and Maley, 2012). The common explanation is that drug therapy often targets the dominant subclone at the time of diagnosis, and upon its remission, either an expansion of previously suppressed subclones, non-susceptible to the treatment, or an emergence of new resistant subclones is likely to happen (Gillies et al.,2012). In the case of monoclonal tumour progression, the temporal order in which specific mutations have occurred has been shown to be informative for disease progression and susceptibility to drug therapy (Ortmann et al., 2015). Therefore, a more comprehensive understanding of the genetic diversity of individual tumours and their evolutionary history is likely to be a key component in the design of more effective personalized cancer therapies (Greaves and Maley, 2012; Stratton et al., 2009; Swanton, 2012).

All cells in a tumour are related via a binary genealogical tree [Fig. 1(b)]. To reconstruct their evolutionary history based on single nucleotide variants, the infinite sites assumption is typically made, which implies that the mutation profiles of the cells [Fig. 1(c)] form a perfect phylogeny A perfect phylogeny exists if for all pairs of mutations *i*_1_, *i*_2_, the set of cells having mutation *i*_1_ and the set of cells having mutation *i*_2_ are either disjoint or one is a subset of the other (Gusfield, 1997). Most approaches to reconstructing tumour phylogenies focus on the partial (temporal) order among the mutation events [Fig. 1(d)]. This tree type implicitly defines the set of possible subclones via the mutation profiles that can be read from the tree by collecting the mutations on the path from the root to any other node in the tree. Not all possible subclones, in particular those at inner nodes, need to have surviving cells. Also by chance cells from surviving subclones may not be sampled.

The main challenge in obtaining knowledge on intra-tumour heterogeneity is that common bulk high-throughput sequencing admixes the DNA of millions of cells in a sample before sequencing. The mutation profiles obtained from the mixture constitute an average of an unknown number of unknown subclones each making up an unknown fraction of the mixture (Navin, 2014). Therefore tree reconstruction needs to be completed by a deconvolution of the mixed signal to identify the subclones, the taxa of the tree. In the past years an abundance of tools have been developed to study subclone composition in mixed samples (Gerstung et al., 2012; Ha et al., 2014; Oesper et al., 2013; Roth et al., 2014; Shah et al., 2012; Zare et al., 2014). Among the approaches that additionally reconstruct the evolutionary relationships, the majority separates subclone estimation and tree reconstruction (Deshwar et al., 2015; El-Kebir et al., 2015; Hajirasouliha et al., 2014; Qiao et al., 2014; Strino et al.,2013), while others combine both tasks into a single step (Jiao et al., 2014; Malikic et al., 2015; Popic et al., 2014). The typical output of these tools would be one or several trees as in Fig. 1(d), augmented with the estimated prevalences of the different subclones in the tumour. Signals from multiple samples from different locations in the tumour increase the statistical power (Deshwar et al., 2015; Jiao et al., 2014; Malikic et al., 2015; Popic et al., 2014). Samples taken from different time points are useful as well (Schuh et al., 2012) but are usually not available for solid tumours, as biopsies are typically taken only once, at the point when the tumour is removed from the patient.

Approaches using mixed samples provide valuable insights into intra-tumour heterogeneity. However, their resolution is inherently limited and inference of both complex subclone structures and low-frequency subclones remains difficult (Navin, 2014; Van Loo and Voet, 2014). The advent of single-nucleus sequencing techniques has started to change the situation. Here, the taxa are known in the form of the individual cells sequenced from a tumour. However, the data we obtain from single-cell sequencing experiments is notoriously error-prone, in particular the false negative rate can be extremely high (≥ 10%) due to the high allelic drop-out rate in the DNA amplification process. The false positive rate is also elevated in comparison to bulk sequencing. Lastly, unobserved sites can be a problem. For example 58% of the data points are reported as missing due to low quality in an early single-nuclei sequencing data set (Hou et al., 2012) thus giving no information on whether the site is mutated or not in the respective cell. This combination of error types prohibits the application of standard perfect phylogeny reconstruction approaches. While generalizations of the perfect phylogeny problem to deal with imperfect data exist, they are typically NP-hard, and modify the input data in the binary mutation matrix, either by finding the minimum number of entries that need to be changed to remove all inconsistencies (Chen et al., 2006), or by removing the minimum number of samples (taxa) to remove all the inconsistencies (Gusfield et al., 2007).

Probabilistic approaches are an alternative to make use of all information contained in the (inconsistent) data. In addition using a Bayesian scheme, the whole posterior tree distribution instead of just a single tree can be obtained and model parameters such as the error rates of the sequencing experiments can be learned. Bayesian approaches typically use polynomial-time Markov chain Monte Carlo sampling heuristics to explore a (super-)exponential search space.

A fully Bayesian approach is BitPhylogeny (Yuan et al., 2015) which uses non-parametric clustering in combination with a tree-structured stick-breaking process to identify subclones and their evolutionary relationships. Unlike tree-based approaches for mixed samples BitPhylogeny clusters samples into subclones and sets these in a phylogenetic relation [Fig. 1(e)].

Kim and Simon (2014) introduced a pairwise ordering test for mutations to attempt to find the best fitting tree from noisy and incomplete single-cell data (Hou et al., 2012). Their approach reconstructs a mutation history as in Fig. 1(d), also referred to as *mutation tree*. The restriction to pairwise tests results in an efficient polynomial time algorithm but comes at the cost of a potential loss in reconstruction quality, as all information from more complex relations than pairwise order is discarded. Instead of using pairwise orders, one could consider testing the ordering of triplets of nodes and then higher groupings.

Here we propose a likelihood-based approach to test the entire mutation tree at once and perform a stochastic search to find the best fitting tree. We introduce SCITE (**S**ingle **C**ell **I**nference of **T**umor **E**volution), a flexible MCMC sampling scheme that allows us to compute the maximum likelihood tree plus attachment points of the samples, sample from their posterior or treat mutation trees with the attachment points marginalised out. These can be combined with learning the error rates of the sequencing experiments. We evaluate SCITE on real cancer data, showing its scalability to present day single-cell sequencing data and its improved results over BitPhylogeny (Yuan et al., 2015), the Kim and Simon (2014) approach, classic perfect phylogeny reconstruction, and methods designed for bulk sequencing data. In addition, we estimate from simulation studies the number of cells necessary for reliable mutation tree reconstruction which could inform the design of future single-cell sequencing projects.

## Results and discussion

### Tree inference from single-cell mutation profiles

We first provide a brief description of our approach to tree inference from single-cell mutation profiles. We start with a model for representing single-cell mutation histories and the likelihood based approach to deal with sequencing errors. Then we give an overview on the different variants of the MCMC sampling scheme implemented in SCITE. A more technical description of SCITE is found in the Methods section.

#### Model of tumour evolution and tree representation

We restrict the evolutionary model to point mutations in this work and make the infinite sites assumption, which states that every genome position mutates at most once in the evolutionary history of a tumour. No further constraints are necessary, in particular no assumption on a mono-clonal origin of the tumour is made, a core assumption in tree reconstruction from mixed samples.

We represent the mutation status of *m* single cells at *n* different loci in a binary *n* × *m mutation matrix E* where a 1, respectively a 0, at entry (*i*, *j*) denotes the presence, respectively the absence, of mutation *i* in cell *j* [Fig. 1(c)]. With the exclusion of convergent evolution due to the infinite sites assumption, this matrix defines a perfect phylogeny of the single cells. This means that there exists a rooted binary tree with the cells as leaves in which every mutation can be placed on one edge such that the mutation status of every leaf equals the set of mutations on its path to the root [Fig. 1(b)]. Mutations present in all cells can be removed from the data as their location in the tree is known. The same is true for mutations observed only in a single cell. These are directly associated with the cell and non-informative in the tree reconstruction. For example, the mutation matrix from Fig. 1(c) reduces to:

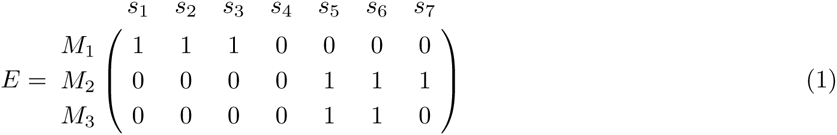

where we now represent the remaining three mutations as *M*_1_, *M*_2_ and *M*_3_. In general, the binary tree defined by the matrix *E* will not be unique. In the example in Fig. 1(b) the three left-most leaves all have the same mutation status, their branching order in the tree is therefore arbitrary. Also the correct placement of the fourth leaf is not unique, as it has no mutation other than the ones shared by all samples. It could equally well branch off in the left subtree after the two ubiquitous mutations instead of the right one. A more compact tree representation of *E* is a *mutation tree T* which represents the mutations as nodes and connects the nodes according to their order in the evolutionary history. An empty node is used to indicate the root [Fig. 1(d)]. The mutation tree can be seen as the perfect phylogeny tree, where instead of placing the mutations along the edges we encapsulate them inside internal nodes. This slight change in representation facilitates our inference later. The mutation tree can be augmented with the sequenced cells by attaching them to the node that matches their mutation state [Fig. 1(f)]. The order of mutations shared by the exact same set of cells is unidentifiable in the mutation tree, as is the case for the two top-most mutations in Fig. 1(f). Such subsets of mutations are summarized in a single node, here highlighted as a shaded box.

### Observational errors

In real data, we do not observe a perfect mutation matrix [Fig. 1(c)] but a noisy version of it [Fig. 1(g)], which we denote by *D* in the following. If the true mutation value is 0, we may observe a 1 with probability *α* (false positive), and if the true mutation value is 1, we may observe a 0 with probability *β* (false negative) such that

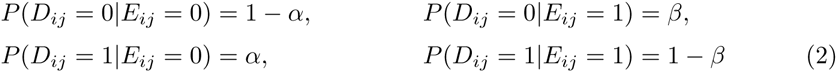

Assuming the observational errors are independent of each other, the likelihood of the data given a mutation tree *T*, knowledge of the attachment of the samples *σ* and the error rates ***θ*** = (*α*,*β*) is then

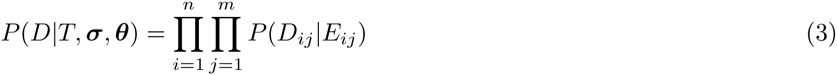

where *E* is the mutation matrix defined by *T* and ***σ***.

For the posterior

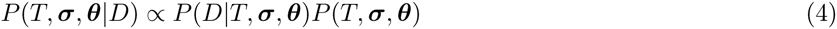

we can factorise the prior, *P* (*T*, ***σ***, ***θ***)= *P*(***σ***|*T*, ***θ***)*P*(*T*, ***θ***), and we assume independence of the error rates to set *P* (*T*, ***σ***, ***θ***)= *P*(***σ***|*T*)*P*(*T*)*P*(***θ***) so that the attachment prior *P*(***σ***|*T*) depends on *T*. Such a prior might be useful if one suspects that cells are more likely to be sampled from later stages in tumour development and lower down in the tree. Here though we use a uniform attachment prior.

### MCMC sampling

Our model for learning mutation histories from single-cell mutation profiles consists of three parts, the mutation tree *T*, the sample attachment vector ***σ*** and the error rates of the sequencing experiment ***θ***. The resulting search space has a continuous component for ***θ*** and a discrete component of size (*n* + 1)^(*n*–1)^(*n* + 1)^*m*^ for (*T*, ***σ***) which prohibits an exhaustive search. Instead, with Equations (3) and (4) we built SCITE, a MCMC scheme to sample from the joint posterior given the data. From the current state (*T*, ***σ***, ***θ***) we propose a new state (*T′*, ***σ**′, ***θ***′*) with an ergodic mixture of moves where we change one component at a time. With properly defined transition probabilities and acceptance ratio our chain converges to the posterior. In practice, we marginalize out the sample attachments in our model not only to speed up convergence but to focus on the mutation tree *T* as the informative part for understanding the mutation history.

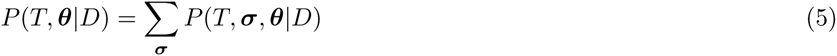

We then only need to consider moves in the joint (*T*, ***θ***) space, thereby reducing the search space by a factor of (*n* + 1)^*m*^. It is still possible to augment the tree with the samples in a post-processing step by sampling them conditionally on the tree.

After convergence, the MCMC chain can be used to sample trees and error rates proportionally to the joint posterior distribution in Equation (4). In addition, it is possible to obtain a single best fitting combination of mutation tree and error rates via point estimates of the model parameters. One way of doing this is via maximum a posteriori probabilities (MAP)

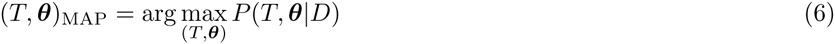

Another possibility is to use maximum likelihood (ML) estimates. Since the likelihood depends on the full set of model parameters (*T*, ***σ***, ***θ***), it is more natural to optimise them all jointly rather than marginalizing out the sample attachment:

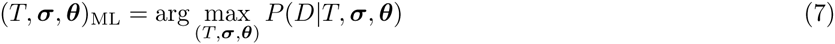

In the ML framework, SCITE includes a parameter γ which amplifies the likelihood and which can speed up discovery of the ML tree.

Finally SCITE provides the option to skip the learning of error rates when fixed error rates are provided. Since these are often available for sequencing data, they can be used instead to reduce the search space size.

## Reconstructing mutation histories from real tumour data

For a first evaluation of SCITE, we applied it to three real single-cell tumour datasets of different data quality.

### JAK2-negative myeloproliferative neoplasm

The first tumour data is single-cell exome sequencing data from a JAK2-negative myeloproliferative neoplasm (essential thrombocythemia) (Hou et al., 2012). It originally consists of 712 SNVs detected in the exomes of 58 tumour cells. In our evaluation we focus on the 18 mutation sites selected as cancer-related by Hou et al. (2012). The error rates of the sequencing were estimated as ***σ*** = 6.04 × 10^‒6^ (false positives) and *β* = 0.4309 (false negatives, allelic dropout). In addition the reduced set has 45% missing data points (compared to 58% in the full data set). The mutation matrix [Supp. Fig. 1(a)] is taken from Kim and Simon (2014). It distinguishes three observed states: normal, heterozygous and homozygous mutation. This only means that a homozygous mutation is observed, not that it is actually present in the data. The latter would contradict the infinite sites model, that each site mutates at most once. Explanations consistent with infinite sites are that we either have a false negative for the normal copy of a heterozygous site, or less likely, a combination of a false positive and an allelic dropout for a site whose true state is homozygous normal. Another explanation for observing a homozygous mutation could be a loss of heterozygosity. We adapted our approach to integrate the third mutation state by using the same error probabilities as Kim and Simon (2014). They assume that the allelic dropout is equally likely to cause a heterozygous mutation to be recorded as a normal state or as homozygous. Denoting heterozygous sites by 1 and homozygous sites by 2, this assumption results in the error probabilities

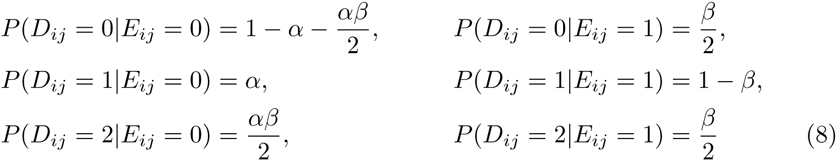

#### Mutation tree reconstruction

We computed the ML tree for the 18 mutation sites with SCITE. When optimising tree and sample attachmet, we obtain a mostly linear mutation tree with a single branching in the lower part of the tree [Supp. Fig. 2(a)] with a ML log score of –378.4.

**Figure 2.**
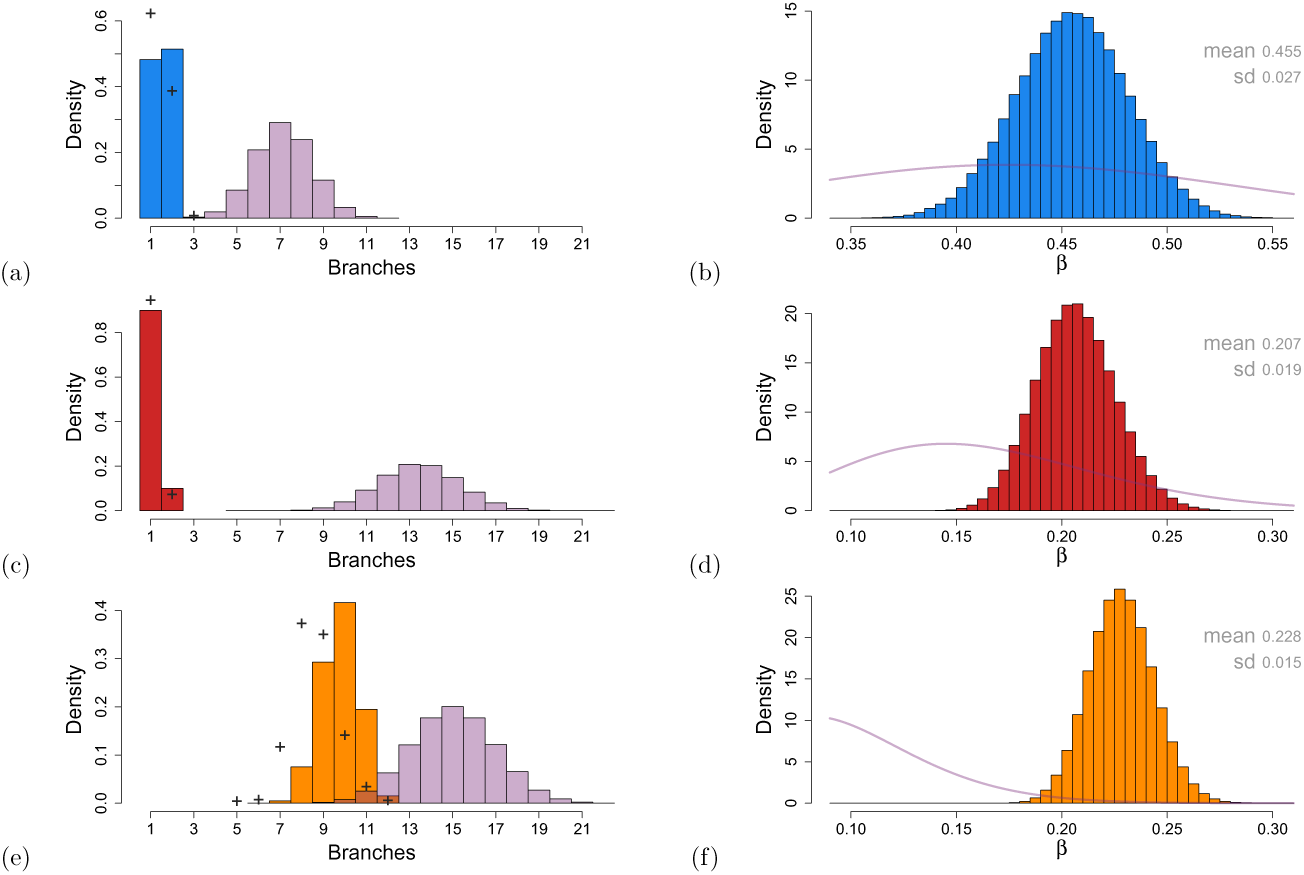
The posterior tree branch and error distributions. The posterior distribution for the number of tree branches for the Hou et al. (2012) data in (a), for the Xu et al. (2012) data in (c) and for the Wang et al. (2014) data in (e), all with fixed false negative error rate *β*. The prior distributions from uniformly sampled trees are in light purple. The posterior distributions for *β* for the same data sets are given in (b), (d) and (f) with the priors included as light purple lines. When *β* is learned, the posterior distribution of the number of tree branches shifts slightly as indicated by the black crosses in (a), (c) and (e).

We observe that quite a few samples are placed at nodes high up in the tree (Supp. Fig. 3) though many of these placements are uncertain, as indicated by the multiple co-optimal attachments. Taking into account the uncertainties due to the high error rates and the large number of missing values (45%), it is not unexpected that many cells fit equally well to several neighbouring nodes. The linear nature of the tree matches a sequential monoclonal development. The subclone expansion starting towards the bottom of the tree indicates the co-existence of multiple subclones at the point of sampling. However, from the single time point data it is not possible to decide whether the more recent subclones are on the verge of replacing the more ancestral clones, or will coexist for longer.

**Figure 3.**
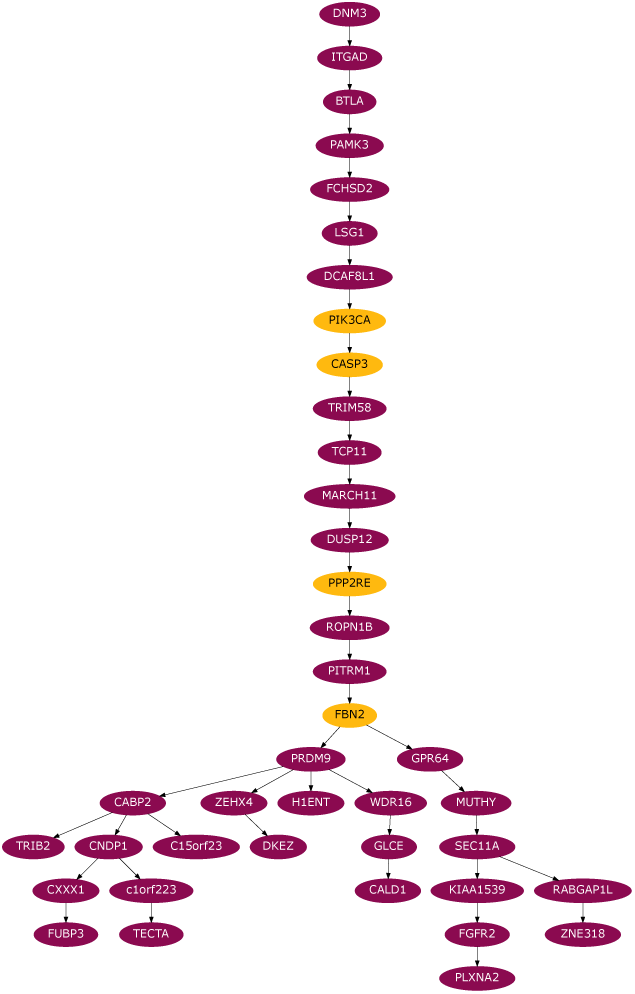
MAP tree for the (ER^‫^) breast cancer. The MAP tree for the Wang et al. (2014) data. See Supp. Fig. 9(a) for a version with samples attached. Yellow genes indicate non-synonymous mutations in known cancer genes (Wang et al., 2014).

Along with finding the ML tree with attachments, we performed a fully Bayesian sampling of trees and attachments from the posterior. To summarise such a sample, we consider as an example the number of branches the trees possesses. The distribution for the Hou et al. (2012) data [Fig. 2(a)] shows that the trees mostly have a single branching point (with two branches) like the ML tree and often occur as a simple linear chain with a single branch.

#### Comparison to trees found with other approaches

The same data has previously been analysed with two competing methods (Kim and Simon, 2014; Yuan et al., 2015).

Kim and Simon (2014) employ the same underlying likelihood with errors as in Equation (8) but they use the data to learn ancestral relations between each pair of mutation nodes instead of the whole tree at once. They also use the data to learn a parameter representing how quickly the mutation tree branches. This parameter is then used to calculate the prior probability of ancestral relations which is fed into their pairwise test and subsequent tree reconstruction.

With the Hou et al. (2012) data (on the same 18 selected mutations) Kim and Simon (2014) estimate that 92% of the evolutionary time of the phylogenetic tree should be before the first binary split. In their model, this translates into expecting over 80% of the mutations to occur before any branching in the mutation tree. Despite this very linear tree estimate, their algorithm to turn the pairwise ancestral relations into a mutation tree leads to the very branched tree in Supp. Fig. 2(c) which has a much lower log-likelihood of –1059.7 than the ML tree found with SCITE (with a log-likelihood of —378.4). This may be due to the use of the minimum spanning tree algorithm by Kim and Simon. The method effectively needs to turn ancestral relations into strict parent-child relations and thereby essentially discounts the deeper history embedded in their pairwise tests.

We cannot compare directly to the tree found by BitPhylogeny (Yuan et al., 2015) since their algorithm aims to find the phylogenetic connection between the samples themselves rather than the mutation tree. Furthermore, the algorithm groups samples into clones according to the data and a stick-breaking prior. For example, using all the mutation data from Hou et al. (2012), as well as a bulk normal and bulk cancer sequence, and with a particular stick-breaking tree prior they find one large clone accounting for over half the samples and 8 further smaller clones arranged in a tree structure (Yuan et al., 2015). However we can view their result as a mutation tree with attachments where the mutations themselves have been censored. This leaves just the sample attachment information as well as the global tree structure between their groupings.

To build a complete mutation tree we allow each mutation to be placed before any one of the clonal groupings of samples (or completely afterwards). For each mutation, we find its ML position and hence find the ML tree (with attachments) which respects the result of Yuan et al. (2015). The resulting tree [Supp. Fig. 2(b)] is a mostly linear chain like the ML tree SCITE finds and involves some of the same genes at the branches although one of our branches is lost. The log-likelihood of –642.3 for this tree is substantially better than the tree of Kim and Simon (2014) but worse than the tree SCITE finds (with a log-likelihood of –378.4). With singlecell sequencing we can, as we do here, simply treat each cell as its own clone and discover their phylogeny directly. BitPhylogeny (Yuan et al., 2015) instead focuses on clustering samples into subclones during tree inference thereby reducing the resolution of the reconstruction.

#### Error rate learning

Within our Bayesian MCMC approach, we can also sample error rates from the posterior. Focusing on the false negative error rate *β* while keeping the false positive *α* fixed, for the beta prior on *β* with mean 0.4309, we chose a large standard deviation of 0.1. In the MCMC chain, with probability 10% a new *β′* is proposed following a Gaussian random walk with standard deviation equal to one third of the prior’s. Running the chain for 10 million steps, throwing away the first quarter and plotting the resulting posterior of *β* we arrive at Fig. 2(b). The posterior mean is 0.455 with standard deviation 0.027 so that the data indicates that the measured value of 0. 4309 is a little underestimated but well within tolerances.

More interesting for our purposes is how these error rates affect the tree inference. The maximum a posteriori (MAP) *β* is 0.455 while the MAP tree (with attachments marginalised out) is a simple chain (Supp. Fig. 4). The mutation order is similar to the ML tree [Supp. Fig. 2(a)] up to the branching point suggesting a monoclonal tumour development. Keeping the error rate fixed at 0.4309 instead, we find an identical MAP tree giving us confidence that the inference is robust against minor differences in the error rates.

**Figure 4.**
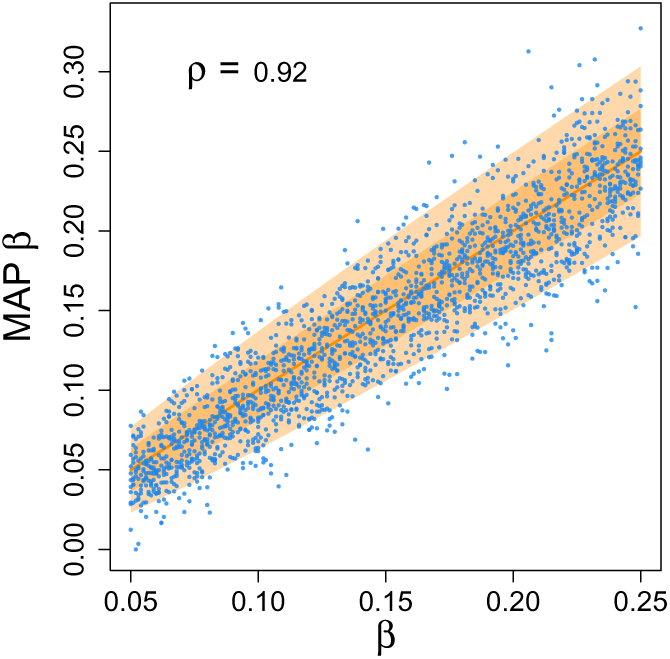
Learning error rates. Comparison of the MAP false negative rate *β* learned using SCITE for *n* =20 against the *β* used to generate the data. The solid blocks are one and two standard deviations of inferring *β* if the tree was known.

#### Mutation tree inference for a larger set of mutations

We also considered a larger set of mutations comprising all 78 non-synonymous mutations from the full dataset. For this number of mutations, with only 58 sampled cells and high levels of missing data (48%), the posterior is rather flat making discovering a global rather than local optimum more difficult. Increasing the parameter *γ* to 2–3 to amplify the likelihood landscape helped in discovering high scoring trees. We also tested that the alternative tree representation [see Methods] designed for instances with more mutations than samples aided in finding the ML tree (Supp. Fig. 5). The ML tree is again highly linear but the order especially of some of the 18 mutations varies compared to the ML tree inferred for that subset of the data (Supp. Fig. 3). With missing data, the mutations may fit equally well along several edges and they were placed in their earliest position which may explain some of the variation. More generally though, the high levels of missing data allow mutations and samples to move without affecting the likelihood while high error rates allow further rearrangements with only a small effect. For example the mutation in the gene PDE4DIP which changes most between the two datasets has 59% missing data. Also the order is essentially determined by the smaller number of samples that attach higher up the trees. This smaller number is effectively reduced further by the missing data limiting the accuracy of any tree reconstruction, as explored later with the Simulations.

**Figure 5.**
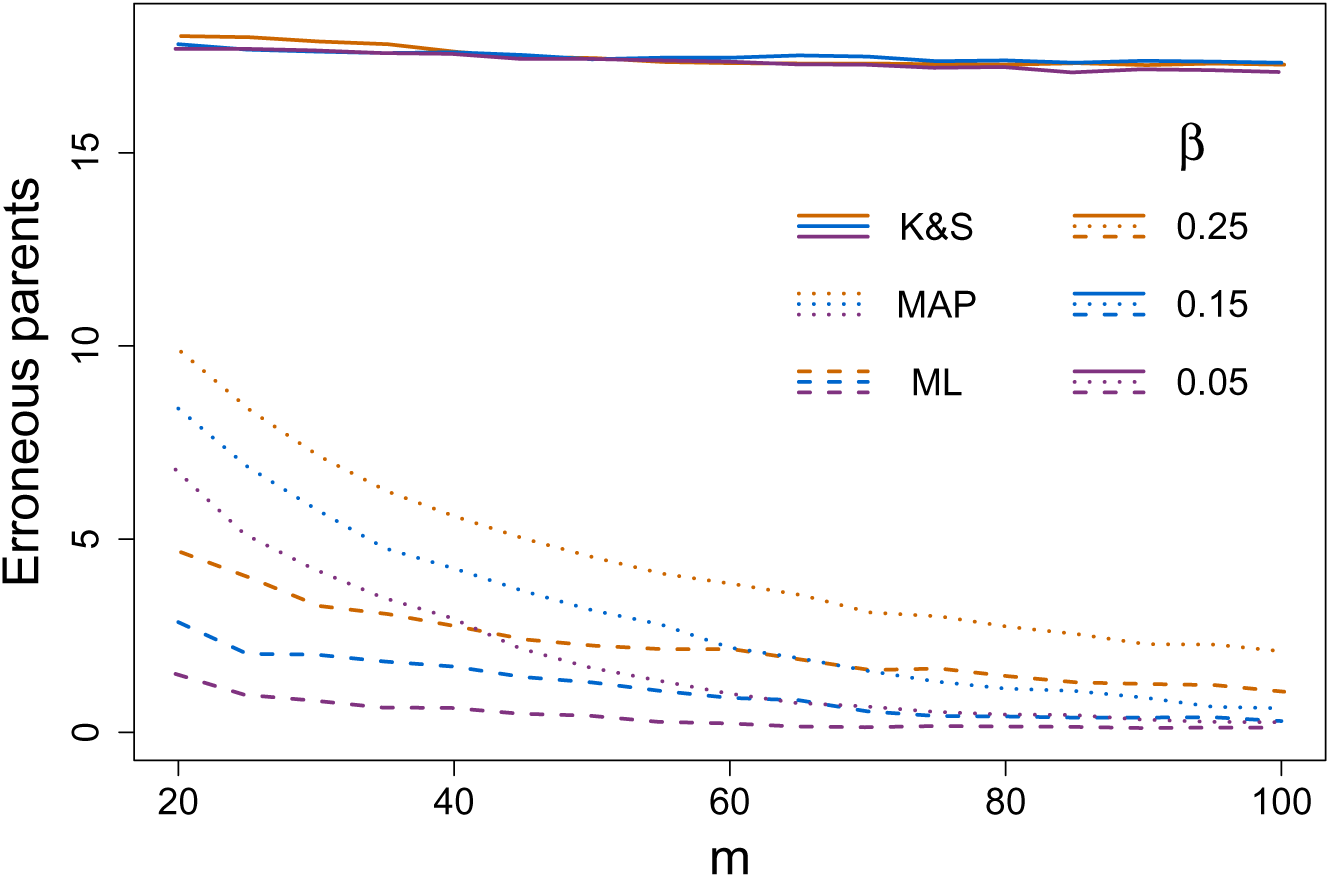
Comparison of different methods. Comparison of the tree learning for *n = 20* using SCITE for the ML tree (dashed) and MAP tree (dotted) against results from Kim and Simon (2014) as solid lines. The ML tree distances do not include non-identifiable regions.

#### Clear cell renal cell carcinoma

The second data set is from single-cell exome sequencing data of a clear cell renal cell carcinoma (Xu et al., 2012). The mutation status of 50 sites in 17 tumour cells are detailed in the supplementary material of Xu et al. (2012) and we marked the presence of an SNV when the call was different from the consensus of 5 normal tissue cells (in line with the totals provided in their supplementary material). As for the Hou et al. (2012) data, Xu et al. (2012) distinguish between heterozygous and homozygous mutations so we again use Equation (8). Of the 50 sites, only 35 were not mutated in at least one cell. Only those were selected since the remaining 15 would simply be placed at the top of the mutation tree. The error rates were estimated by Xu et al. (2012) as ***σ*** = 2.67 × 10^‒5^ (false positives) and *β* = 0.1643 (false negatives) and the data also has 22% missing entries [Supp. Fig. 1(b)].

#### Mutation tree reconstruction

The ML and MAP trees both possess a completely linear accumulation of mutations [Supp. Figs. 6 and 7(a)] which is consistent with a series of monoclonal expansions and the conclusions of Xu et al. (2012). The linearity is confirmed in the full posterior distribution of trees with a linear chain being dominant [Fig. 2(c)]. In addition, we observe that almost all of the samples are placed towards the end of the tree. Again a larger value of the parameter *γ* and the alternative tree representation sped up discovery of ML trees.

**Figure 6.**
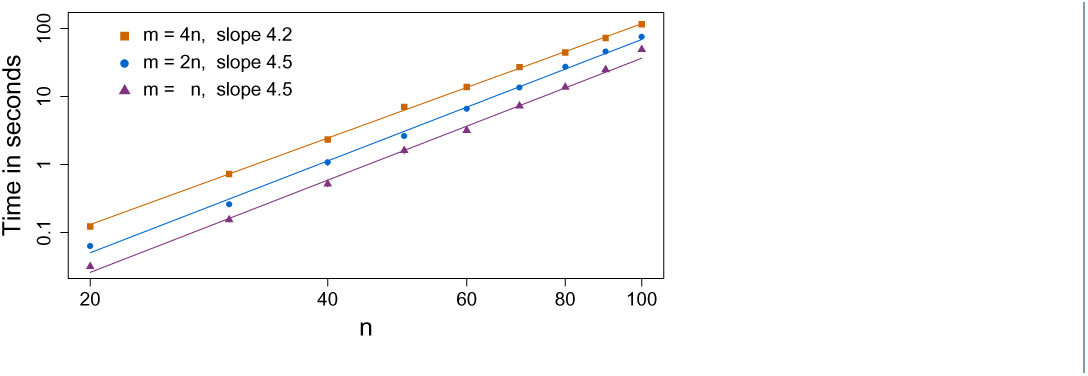
Scaling behaviour. The average time taken for SCITE to first find a ML tree as the number of mutations *n* in the tree is varied along with the number of attached samples *m*={*n*, 2*n*, 4*n*}.

**Figure 7.**
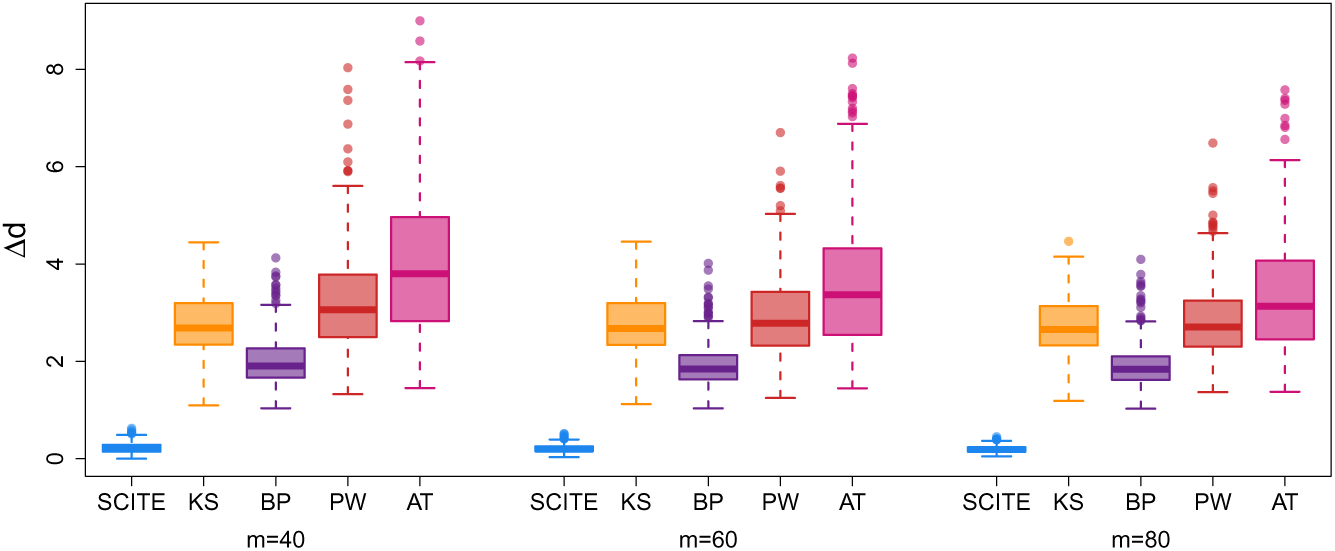
Comparison of additional methods. Comparison of the tree inference of SCITE, the algorithm of Kim and Simon (2014), BitPhylogeny (Yuan et al., 2015), PhyloWGS (Deshwar et al., 2015) and AncesTree (El-Kebir et al., 2015). The quantity Δd is the normalised consensus node-based shortest path distance (as defined in Yuan et al., 2015) between the inferred and generating trees.

#### Error rate learning

Fixing a beta prior for *β* with mean 0.1643 and standard deviation of 0.06 the posterior distribution of *β* was obtained by averaging over 10 runs of 10 millions steps (with a quarter as burn-in) [Fig. 2(d)]. The posterior mean is a little larger at 0.207 with a standard deviation of 0.019 so the stated value is just within the uncertainties. The MAP value of *β* instead is a little closer at 0.198 while the MAP tree [Supp. Fig. 7(b)] is essentially identical to the that with a fixed value of *β* = 0.1643 [Supp. Fig. 7(a)]. The order of some of the higher mutations varies however since their exact placement hardly affects the posterior probability.

#### Oestrogen-receptor positive (ER+)breast cancer

The third dataset is from single-nuclei exome sequencing of 47 tumour cells from an oestrogen-receptor positive (ER^‫^) breast cancer (Wang et al., 2014). Only two states are called for each site, presence or absence of a SNV. Estimated error rates from Wang et al. (2014) are 9.72% for allelic dropout, and 1.24 × 10^‒6^ for false discovery. In our analysis we use the 40 mutations present in at least two tumour cells [Supp. Fig. 1(c)].

#### Mutation tree reconstruction

The MAP tree computed for this dataset is shown in Fig. 3. In the Supplement we additionally show the ML tree [Supp. Fig. 8] and a version of the MAP tree with attached samples [Supp. Fig. 9(a)]. In both the MAP and the ML tree we observe a linear accumulation of mutations in the early stages of the tumour suggesting that the development was through a sequential replacement of subclones with no surviving side branches and only a few cells with ancestral states surviving until present. In the later stages of the tumour, we observe a complex branching into co-existing subclones. This branching is exhibited more generally in the full posterior distribution of trees as summarised in Fig. 2(e).

**Figure 8.**
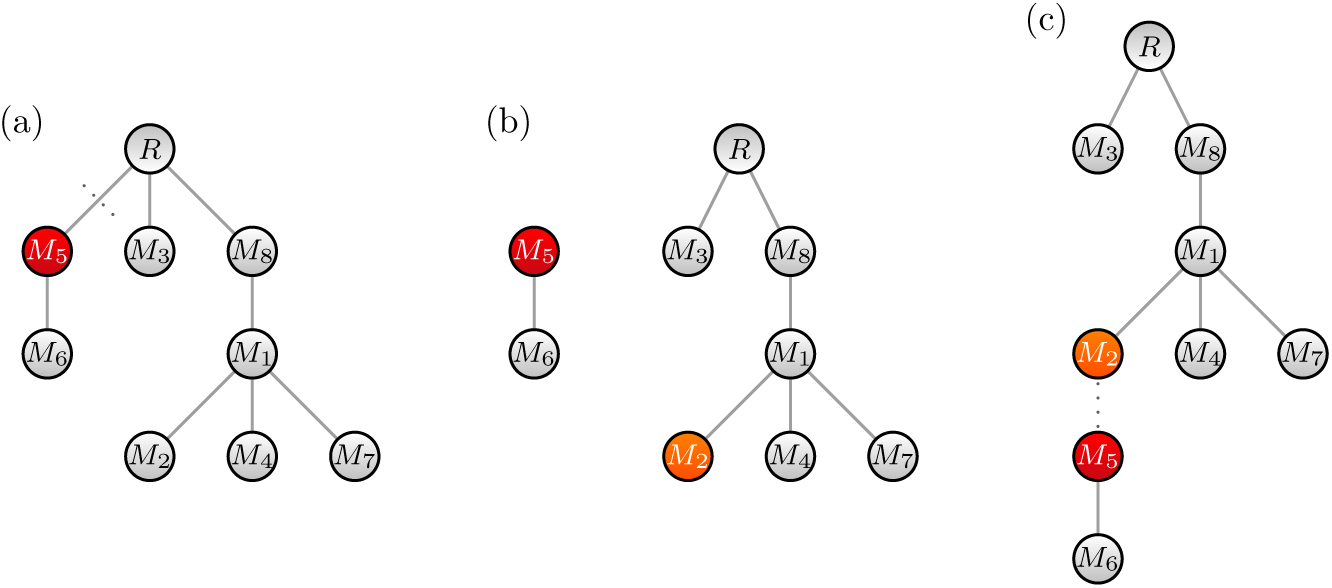
Prune and reattach MCMC move. (a) From our starting tree *T* we first select a node uniformly, here *M*_5_, and detach it from the rest of the tree. (b) Then we sample one of the remaining nodes in the rooted tree, here *M*_2_ is chosen. (c) Finally we attach the detached subtree formed of *M*_5_ and its descendants to the newly selected node *M*_2_.

From the single time point data available for this tumour it can not be inferred whether there will be a long-term coexistence of subclones, or if we observe a transient state which will eventually lead to a single surviving subclone. For initial cancer treatment however the *status quo*, which mutations co-occur in cells, is already informative to jointly target the present subclones and therefore minimise the risk of further differentiation into therapy-resistant subclones.

#### Error rate learning

Using a beta prior for *β* with mean 0.0972 and standard deviation of 0.04 we averaged over 20 runs of 10 million steps (with a quarter as burn-in) to obtain the posterior distribution of *β* [Fig. 2(f)]. The posterior mean is more than double at 0.228 (with a standard deviation of 0.015) which is in disagreement to the stated value. This result is in contrast to our later simulations on learning the error rate [Fig. 4] that show that the MAP value is close to the true one. A possible explanation for the discrepancy is that allelic dropout only comprises one part of the false negative rate. Other contributing factors could include inaccuracies in calling heterozygous mutations at low coverage.

The MAP value of *β* is 0.226 with a MAP tree [Supp. Fig. 9(b)] shares many feature with the MAP tree at fixed *β* = 0.0972 [Supp. Fig. 9(a)] but has some rearrangements of the branches lower down and some reordering of the mutations higher up. Learning the error rate also leads to slightly fewer branches in the posterior distribution, as indicated by the black crosses in Fig. 2(e).

### Systematic evaluation of SCITE on simulated data

With the limited availability of single-cell sequencing data at this point and the lack of the ground-truth in real data, we performed a more systematic evaluation of SCITE on simulated datasets. Our analysis focuses on the accuracy of tree inference and error rate learning, the effect of the data quality, and the practical runtimes of SCITE.

#### Accuracy of tree inference

To check the consistency of our approach, we simulated random mutation trees with attachments uniformly, which allows for poly-clonal tree topologies. First, for *n* = 20 and ***σ*** = 10^‒5^, we generated 100 such trees with up to 100 attachments. For error rates 100*β* ∈ {5,15, 25}, for each tree we sampled from a lognormal with standard deviation 0.1 and multiplied it by *β* to obtain *β*^*^. Then we added noise to the perfect data with rates (*σ*, *β*^*^) and removed 1% of the data. Taking subsets of the data of size *m* we learned the ML and MAP tree for the error rates *β*. This gives us a random misspecification of around 10% compared to *β*^*^.

We quantified the difference between the inferred trees and the true tree by counting how often a node has the wrong parent (Fig. 5 and the top row of Supp. Fig. 10). In the ML setting, if no samples are attached to a chain of mutations, then any ordering of those mutations has the same likelihood. Here, in the score we do not penalise this non-identifiability and take the ordering which minimises the distance to the generating tree. The non-identifiability will however tend to decrease as the number of samples *m* increases. The MAP tree does select an ordering (roughly following the frequencies) and hence has higher distances than the ML tree. In general MAP inference should be more robust and less prone to overfitting, but can have a higher bias. To fairly compare the ML and MAP inference we chose a random ordering of the mutations in non-identifiable regions in the ML trees and recomputed the distances to the generating tree. We do observe a marginal improvement in the tree reconstruction with the MAP tree (Supp. Fig. 11).

The errors however are not a result of the inference method, since SCITE indeed finds the ML tree (Supp. Fig. 12). Instead these errors are inherent in noisy data where another tree might happen to fit the data better than the generating tree. The discrepancy can only be resolved by reducing errors or increasing the sample size and Supp. Fig. 10 gives an indication of how this occurs. To put the errors in scale, a value of two would refer to adjacent mutations in a chain being swapped. Since samples contain the mutations along their entire history in the mutation tree, we have a greater consensus about the mutation structure higher up the tree than lower down. The exact placement of mutations near the bottom of the tree may be determined by only a couple of samples so that the errors we typically see with larger *m* are mutations near the bottom of the tree being shifted, or two adjacent mutations being swapped. With this in mind, we obtain very good trees with about 60 samples, depending on the error rate.

We repeated the simulations for *n* = 40 and up to 200 attachments as depicted along the bottom row of Supp. Fig. 10 and again find good reconstruction when we have several samples per mutation.

#### Learning the error rates

Since SCITE can also perform fully Bayesian tree inference, we examined its ability to infer the false negative rate from data. For 2000 random trees with 60 attachments we generated data with a range of *β* from 5% to 25%,*σ* = 10^‒5^ and 1% missing data. We further fixed a uniform prior for learning *β* so that no information is passed to SCITE apart from the noisy and incomplete mutation matrix.

There is very high correlation between the generating *β* and the MAP value learned (Fig. 4). To put this in context, we consider the theoretical distribution if the tree was known. From the random trees and attachments, around 22% of the entries in the perfect mutation matrix are ones. They are randomly changed with the rate *β* leading to a binomial distribution and a standard deviation of 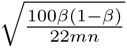 when inferring *β* from the result. One and two standard deviation intervals are included in Fig. 4, showing again that SCITE performs very well as it must also infer the tree structure and handle the missing data.

Similar plots for *m* = 40 and *m* = 80 (Supp. Fig. 13) show also a tightening of the *β* inference as *m* increases.

#### The effect of missing data

High rates of missing data points due to unobserved mutation states are typical for present-day single-cell sequencing data. We performed simulation experiments to test how this feature affects the accuracy of mutation tree reconstruction. With an error rate of *β* = 10% and the same misspecification as before we generated up to 400 random trees with up to 80 attachments. Keeping *σ* = 10^‒5^ we varied the amount of missing data from 1% to 20% to see the effect on the tree reconstruction for *m* = {40, 60, 80}. We see a very weak increase in reconstruction errors as the missing data rate increases (top row of Supp. Fig. 14). Since SCITE treats the inference probabilistically, missing data is akin to effectively reducing the number of samples m, so the behaviour in Supp. Fig. 14 is in line with changing *m* slightly in Supp. Fig. 10. The behaviour also shows that SCITE is robust even against high missing data rates.

Looking back to the even higher missing data rates in the earliest datasets, we simulated up to 60% missing data with 400 trees and the same settings as before. The reconstruction progressively gets worse with increasing missing data (bottom row of Supp. Fig. 14). At around 30-40% missing data with 80 attachments we have similar performance as for 40 cells attached with no missing data, and so have effectively halved our sample size. With 60% missing data, the reconstruction is notably poorer again, although SCITE does find about half the parents correctly for the MAP solution and a large majority with the ML approach. This difference is because the optimal order is chosen for the ML solutions in case of non-identifiablility.

#### Doublet samples

Rarely, instead of isolating a single cell for sequencing, a pair of cells is captured instead. We checked how robust SCITE is to these sorts of perturbations by again simulating data from 400 random trees with 20 nodes and up to 100 attachments. To represent the sequences of doublet samples, we took up to 20 pairs of attached samples and combined them by recording a mutation whenever it was present in either of the original single cells. Errors were added with a rate of *β* = 10% (misspecified as previously), *α* = 10^‒5^ and 1% missing data. We ran SCITE with *m* = {40, 60, 80} total samples, including up to 20 doublets, to see their effect on the tree reconstruction.

We observe a linear increase in reconstruction errors as the number of doublets increases (Supp. Fig. 15) with decreasing gradient as *m* increases since then the doublets represent a smaller proportion of the total sample. Unlike missing data which reduces the effective sample size, doublets add confounding mutations which could disagree with the tree topology. However since SCITE employs probabilistic inference, and at the level of the mutation tree rather than sample tree, the consensus of the single cell samples moderates the negative effects of the doublets. Even at high rates of doublet sampling, like 10 or 20%, the tree reconstruction therefore performs well.

#### Runtimes

To uncover the complexity of the stochastic search and MCMC scheme, we simulated data from 400 uniformly sampled trees with up to 100 nodes and 400 attached samples. We set *α* = 10^‒5^ and *β* = 0.1 (with the same misspecification as before), included 1% missing data and set the parameter *γ* = 1 as for the MCMC case. For each tree we ran SCITE 100 times and recorded how many steps the algorithm took to first hit the highest likelihood tree uncovered by that run, as well as the time of the run. The lengths of the chains was chosen so that nearly all of the runs would share the same highest likelihood. The average number of steps needed to first find the consensus ML tree can then be calculated (for those runs with a lower likelihood we add the length of the chain and then assumed they would find the ML tree in an additional average number of steps). This can then be multiplied by the average time per step to give a measure of how long SCITE takes to find a ML tree on average, and repeated for all 400 trees to provide Fig. 6.

On the theoretical side, arguments analogous to those in Kuipers and Moffa (2015) indicate that the MCMC chain requires *O*(*n*^2^ ln(*n*)) steps to converge or find ML trees. The likelihood landscape may also depend on *n* and *m* in non-trivial ways which can further affect the convergence. With each MCMC step taking O(mn) to score the tree we get an overall estimate of *O*(*mn*^3^ ln(*n*)) for convergence.

Comparing to the numerical results in Fig. 6, the gradients in the log-log plots are 4.5, 4.5 and 4.2 for *m* = {*n*, 2*n*, 4*n*} respectively. Since *m* ~ *n* in the simulations, these are a little higher than the power of 4 suggested by the estimate, but roughly in line with it. To check the linear scaling with m, we take the fit lines at *n* = 60 in the middle of the simulation we find that doubling *m* from *n* to 2*n* and then 4*n* increases the time by a factor of 1.9 and then 1.95, slightly less than double and in line with linear scaling. With linear scaling in *m*, and for a reasonable number of mutations, SCITE will therefore be able to efficiently handle large numbers of sampled cells.

Further parameters with influence on the practical performance of SCITE are the move probabilities, and for ML tree discovery additionally the parameter *γ*. We performed a systematic search for the optimal parameters which is described in the Supplementary Material. Our observation is that an optimal choice of move probabilities gives a constant factor speed-up compared to default values. Similar results were observed for *γ* which has its optimum for finding a ML tree quickly just below 1, the value required for the MCMC sampling.

### Comparison to competing approaches

To further assess the performance of SCITE we compared it to a simple perfect phylogeny approach, two methods designed for single-cell data, and two recent methods for tree inference from bulk sequencing data.

#### Perfect phylogeny

We first compared SCITE against a simple algorithm for solving the perfect phylogeny problem (i.e. testing whether the data defines a phylogeny, and if it does to construct one, Gusfield, 1997). A mutation matrix has a perfect phylogeny if a tree can be constructed such that the leaves are the samples and the mutations are each placed at exactly one edge, such that for every leaf the mutations on the path leading to it from the root reflect its mutation status. Such a tree only exists if there are no contradictions in the data due to noise or recurring mutations. But if it exists, it can be represented as a mutation tree by labelling nodes instead of edges. To test for perfect phylogeny we use a version of the data with no missing values. From our simulated trees and data, only very few are contradiction free which limits the tree comparison to a few instances. The perfect phylogenies on average deviate more from the true tree than both ML and MAP trees and none is found for instances with more than 45 samples. The differences between the perfect phylogeny and the true tree are due to both the errors introduced and insufficient information to fully reconstruct the tree. Details of the comparison are given in Supp. Table 1.

#### The approach of Kim and Simon (2014)

The method in Kim and Simon (2014) reconstructs the same type of mutation trees as our approach. However in their approach, a parameter representing how quickly the mutation tree branches is first learned from the data. This parameter is then used to calculate the prior probability of ancestral relations which informs a pairwise ordering test and subsequent tree reconstruction. Instead of learning the parameter from the data, we give their method the exact value from the tree that was actually used to generate the data since this simplifies running the simulation test. Of course, in practice this piece of information would not be available so the results from their algorithm are over optimistic. Nevertheless, the pairwise approximation performs comparatively poorly [Fig. 5]. In particular, there is little improvement as the number of samples increases. Although the pairwise ancestral tests will become more accurate, this additional information appears to have little impact on the conversion to a mutation tree.

#### Comparison with BitPhylogeny

More advanced probabilistic inference is provided by BitPhylogeny (Yuan et al., 2015). This method, however, reconstructs a hierarchical subclone structure rather than a mutation tree thereby precluding a direct comparison to SCITE and the approach of Kim and Simon (2014). Therefore, we convert the outcome of each method into a complete mutation tree with samples attached. For SCITE this means finding the ML tree with attachments and for the approach of Kim and Simon (2014) we place the samples at their best fitting position on the tree found. For BitPhylogeny instead, we place the mutations along the branches of their clonal tree in the position which maximises the likelihood. Since the mutations and samples may be grouped together, as a measure of fit we use the consensus node-based shortest path distance (as defined in Yuan et al., 2015) between the (completed) inferred tree and the generating tree. In particular, for each tree, the pairwise shortest distance between any two samples is their number of differing mutations. We then normalise by averaging over the absolute differences between the pairwise distances in the inferred and generating trees, rather than taking the sum.

For *n* = 20, *α* = 10^‒5^ and *β* = 0.1 (with the same misspecification as before), we generated 400 such trees with 1% missing data. For simplicity and giving BitPhylogeny a slight advantage, we passed it the complete data. The results for *m* ∈ {40, 60, 80} are presented in Fig. 7. The compared methods perform significantly poorer than SCITE, with BitPhylogeny (Yuan et al., 2015) performing better than the algorithm of Kim and Simon (2014), but with neither approaching the performance of SCITE.

We can also compare the performance of the different methods in terms of the difference in log-likelihoods between the inferred and generating trees, normalised by dividing by the number of data matrix elements (Supp. Fig. 12). This shows similar behaviour to Fig. 7 and we observe that SCITE always provides a nonnegative difference. SCITE therefore always found either the generating tree or one with a slightly higher likelihood than the generating tree.

#### Comparison with bulk-sequencing methods

Finally we compared SCITE to methods designed for deconvolution and tree reconstruction from mixed bulk sequences. We chose PhyloWGS (Deshwar et al., 2015) and AncesTree (El-Kebir et al., 2015) as two recent high-performing methods which allow the samples to be treated separately as well as combined. PhyloWGS employs a stick-breaking tree prior (like BitPhylogeny) while AncesTree solves the deconvolution and ancestry as a matrix factorisation. When passing the simulated single-cell mutations as individual samples to both methods, neither returned anything other than a single grouping of mutations. A possible explanation for this result is that the two methods interpret the binary mutation states as cellular prevalences in mixed samples which likely causes trouble in the deconvolution step. Better performance was obtained when combining the single cells into a bulk mixture, with both methods returning mutation trees with the mutations possibly grouped together at the nodes. To compare with the other methods, we again placed the samples at their best positions in the inferred trees to obtain the results in Fig. 7. AncesTree performs slightly worse than PhyloWGS and both are notably worse than BitPhylogeny and SCITE. This is not unexpected, as only the latter two are designed to handle single-cell data. The main conclusion here is that specialized methods are necessary for single-cell data as approaches for mixed samples are not readily applicable.

## Conclusions

Single-cell sequencing data is giving unprecedented insights into intra-tumour heterogeneity, a major obstacle to permanent remission in cancer treatment. In this paper we introduced SCITE, a likelihood-based reconstruction of tumour genealogies from noisy and incomplete mutation profiles of single cells. The approach provides a flexible MCMC sampling scheme that allows to either find the best fitting tree, or sample from the posterior distribution, and can be combined with learning the error rates of the sequencing experiments. We have shown that the probability model underlying SCITE is highly adaptable. It performs well in the presence of various types of noise, including types that were not explicitly modelled, such as doublet samples (the inadvertent sequencing of two instead of a single cell). The model also lends itself to some straight-forward extensions such as the incorporation of position specific error rates, or the introduction of further mutation and error types that would maintain the independence of genome positions. Besides its flexibility, the key advantage of SCITE is its linear scaling with the number of samples. While this feature is negligible for present datasets, it will become essential as soon as hundreds or even thousands of cells of a tumour will be routinely sequenced.

Using SCITE we reconstructed the mono-clonal origin of an ET tumour and a clear cell renal cell carcinoma, as well as a complex sub-clonal structure in an ER^‫^ breast cancer. The consistency of SCITE is shown in simulation studies which we also used to estimate the number of cells necessary to obtain reliable tree reconstructions, a piece of information that could be useful in the design of future sequencing experiments.

SCITE differs from earlier approaches, in particular BitPhylogeny (Yuan et al., 2015), in its use of single cells as taxonomic units giving it the highest possible resolution in the tree reconstruction. Because each cell provides information about all the mutations, and all this detail is used, this approach allows a more robust reconstruction of the mutation tree. This in turn aids the identification of driver mutations. The placement of the individual cells is however less certain. Clustering cells into clones instead, as done in BitPhylogeny (Yuan et al., 2015), and placing these as the taxa means we can use the consensus of single-cell information in each clone to more robustly deduce the ancestral relationships between the clones themselves, but at the expense of reduced accuracy in the reconstruction of the mutational history.

Further improvements in tree reconstruction could be achieved by considering copy number alterations along with point mutations. For one, copy number information could be used to better understand point mutations states, e.g. a seemingly homozygous mutation may in fact be loss of heterozygosity, but more importantly it can be used as a feature in tree reconstruction itself, as has been done previously for bulk-sequencing data (Deshwar et al., 2015). The main challenges here will be that for large-scale copy number events independence of mutation sites is no longer given, and that the infinite sites assumption would no longer hold.

The knowledge of individual mutation histories is a promising source of information for personalized cancer treatment. Once single-cell sequencing has become more prevalent, the unprecedented resolution of mutation histories reconstructed from single cells will likely be valuable in many more respects. One direction is the identification of recurrent mutation patterns by comparing high-resolution mutation trees from patients with the same and/or different tumour types. Another direction could be to combine single-cell data from different time points and different locations in the tumour to obtain a better understanding of the temporal and spatial organisation of sub-clonal populations of tumour cells, again at a higher resolution than would be possible with bulk sequencing data. When sampling cells from the primary tumour and metastasis, the attachment point of metastatic cells to the mutation tree could help to answer the open question whether subclones with the potential to metastasise arise early or late in the tumour development.

## Methods

### Mutation trees

In SCITE we represent a rooted mutation tree *T* over *n* mutations as an augmented ancestor matrix *A*(*T*) where every node is considered an ancestor of itself:

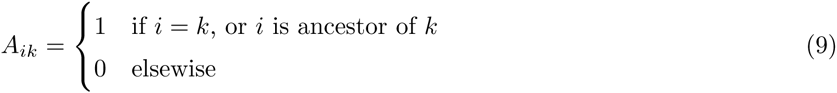

For example the augmented ancestor matrix for the tree in Fig. 1(d), reduced to the mutation matrix given in Equation (1) is

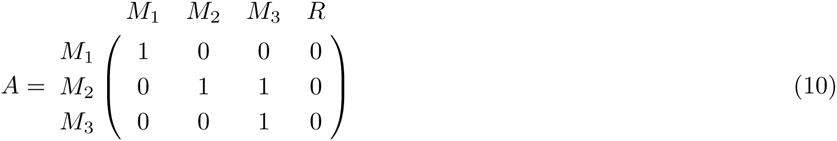

where *R* represents the root of the mutation tree. The cells are attached to *T* such that the path to the root spells out their mutation status. This placement is denoted by a vector ***σ*** which records at the *j*-th position the attachment point of sample j. Carrying on the example from 1 and Fig. 1(d), we have ***σ*** = (1,1,1,4, 3, 3, 2) where 4 represents the root. The connection between the mutation matrix and the mutation tree is

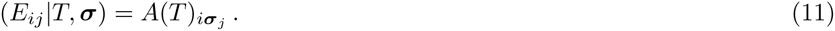

This simply means that for a given tree and sample attachment, the mutation status of a sample is identical to the one observed in the node where the sample attaches to the tree. Therefore the likelihood in Equation 3 can be rewritten as

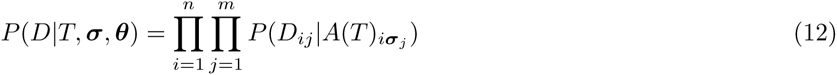

and thereby computed directly from *T* and *α*.

### MCMC sampling

In principle, the MCMC of SCITE needs three types of moves to separately alter the tree *T*, the attachment vector ***σ*** and the error rates. In fact, it is possible to marginalize out the ***σ*** component, such that we only need to consider moves in the joint (*T*, ***θ***) space. We first focus on the marginalization and then describe the remaining move types.

### Marginalization of the sample attachment

A move where we pick a sample and a new parent for it uniformly would satisfy the necessary properties for the MCMC chain on ***σ*** to converge, but we can achieve convergence much faster. This is because the likelihood in Equation (12) factorises into a product for each sample to be attached. As long as the prior *P*(***σ***|*T*, ***θ***) can also be factorised (so that the attachment for each sample is independent of the others) we can include the priors as in Equation (4) and efficiently sum Equation(12) over ***σ*** to marginalise it out

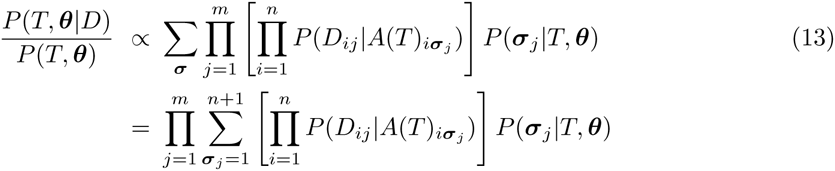

Computation of Equation (13) is in *O*(*mn*) time due to the tree structure underlying *A*(*T*). Along with efficient computation of Equation (13) we now need only search over the (*n* + 1)^*m*^ times smaller space of trees *T* and error rates ***θ*** leading to much faster MCMC convergence. This marginalisation is equivalent to grouping all attachments to the same tree into a single object which is the idea responsible for the similarly large speed up for sampling Bayesian networks in order MCMC (Friedman and Koller, 2003) and more recently partition MCMC (Kuipers and Moffa, 2016). Analogously Nested Effect Models average over all effects with uniform prior (Markowetz et al., 2007).

With the attachments marginalised out, we only need to consider moves in the joint (*T*, ***θ***) space. We can change one component at a time to propose a new pair (*T*′, ***θ***′) with transition probabilities *q*(*T*′, ***θ***′|*T*, ***θ***) and accepting moves with the ratio

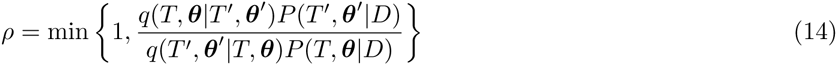

to sample proportionally to *P*(*T*, ***θ***|*D*). Once we have sampled a tree we can easily sample each attachment independently following Equation (13).

#### Tree moves

Akin to standard MCMC approaches on graphical structures (Madigan and York, 1995) we build a scheme on rooted mutation trees for fixed errors ***θ*** as follows. Given the current tree *T* we find the neighbourhood of all trees reachable with the MCMC move from *T*. One then samples a tree *T*′ from this neighbourhood with some proposal probability q(*T*′, ***θ***|*T*, ***θ***) and accepts the move with the probability in Equation (14).

As long as the moves satisfy reversibility (that is, if the move from *T* to *T*′ can be proposed with a non-zero probability, the reverse move from *T*′ to *T* also has non-zero probability to be proposed), irreducibility (that is, a sequence of moves exists that leads from any tree to any other) and aperiodicity (which can be ensured by including the tree *T* in its neighbourhood or adding a non-zero probability not to move), once the chain converges this scheme would allow us to sample trees proportionally to *P*(*T*, ***θ***|*D*).

The basic MCMC move we use is *prune and reattach*. We sample a node *i* uniformly from the *n* available and cut the edge leading to this node to remove the subtree from the tree. Then we sample one of the remaining nodes (including the root) uniformly and attach the subtree there instead. An example is illustrated in Fig. 8.

The reverse move, where we again sample *i* first but then pick its old parent, has the same proposal probability *q*(*T*, ***θ*** |*T*′, ***θ***) = *q*(*T*′, ***θ***|*T*, ***θ***) since the non-descendant set has the same size each time *i* is removed. This term then drops from Equation(14) and need not be calculated. Since we can also choose the old parent when sampling a new one, this move has a non-zero probability of proposing the same tree *T* ensuring aperiodicity. There is also a path from any tree to a tree with all nodes attached to the root, by moving each node to the root step by step. Via reversibility we can likewise move from there to any other tree ensuring irreducibility. The *prune and reattach* move therefore suffices to sample trees according to their posterior. To speed up the convergence of the chain, we use two additional moves in our MCMC scheme: *swap nodes* to swap the labels of two nodes and *swap subtrees* to swap two subtrees. (See the Supplementary Material for more details.) One of the three moves is picked at each step of the chain with a fixed probability.

#### Error learning

Where estimates of the error rates are known we can input this information into the prior *P*(***θ***). Since ***θ*** is between 0 and 1 we choose a beta prior with mean equal to the known estimates and a large standard deviation to be weakly informative. Although for a given (*T*, ***σ***) we can marginalise out ***θ*** analytically, this interferes with the speed up in Equation (13). Instead to move in the error space we choose a simple Gaussian random walk with fixed standard deviations in each direction and centred on the current value ***θ***.

#### Maximum likelihood tree

Along with utilising our MCMC scheme to perform fully Bayesian tree inference, we can adapt the method to search for the ML tree as well. In the ML framework we consider the full space of trees with attachments (*T*, ***σ***) and find the best scoring pair, rather than summing over the attachment points. Keeping the error rates ***θ*** fixed for simplicity, when we wish to search for the ML tree with attachments, we can define the following score for each tree

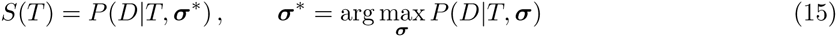

after maximising over all possible placements. Since the likelihood in Equation (12) factorises we can find the best attachment for each sample independently of the others

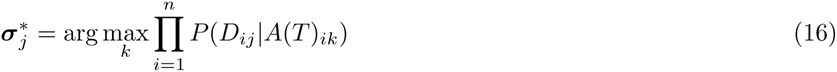

by running over the columns of *A*(*T*) and comparing to the observed data with the error rates as in Equation (2). If several placements provide the same maximum, any may be selected for calculating *S*(*T*) which is then

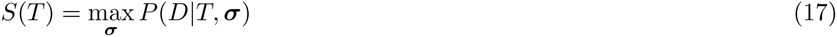

and which again involves only *O*(*mn*) simple operations.

Now we can turn our attention to finding the ML tree

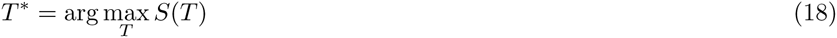

which because of Equation (17) is the tree which maximises the likelihood in Equation (12). The number of rooted trees with *n* + 1 nodes (including the root) grows factorially so an exhaustive search becomes infeasible for more than 10 nodes or so.

Instead we can reuse our MCMC scheme on the space of rooted mutation trees where given the current tree *T* we propose a tree *T*′ according to one of the three move types with the same proposal probability *q*(*T*′ |*T*) but now accept the move with probability

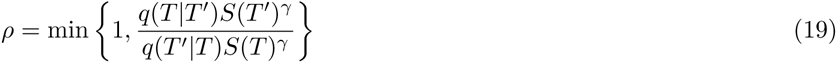

The power of *γ* here is a way to flatten the distribution (for *γ* < 1) or make it more pronounced (for *γ* > 1). Simulated annealing would involve running the chain while simultaneously increasing *γ* → ∞ to to end up in a local maximum. Here instead we chose a value (depending on the data) for this parameter which allows fast discovery of the maximally scoring tree and simply run many chains recording the maximally scoring tree we encounter.

In the Bayesian framework, we can search for the MAP tree. Including the prior on all discrete components and updating *S*(*T*) accordingly we would find the joint MAP tree and attachments with the scheme here. Averaging out the attachments instead we can search just for the MAP tree as well. In particular we replace *S*(*T*) by *P*(*T*, ***θ***|*D*) in Equation (19). We can also find jointly maximal trees and error rates by putting the error moves back in.

#### Alternative representation for ML discovery

For the ML tree with attachments, since the optimal placement for each attachment can be easily found, we are left to search over all rooted trees with (*n* ‫1) nodes. However when *m* ≲ *n* we may return to the binary genealogical tree [Fig. 1(b)] with *m* sampled cells as leaves and (*m* – 1) internal binary divides. The mutations are placed along the edges and present in all cells further down that lineage. For a given genealogical tree with leaves, the optimal placement of every mutation along the edges is simple to compute. Each tree is assigned a score corresponding to the likelihood of the data given the optimal placement of the mutations, which can again be calculated in *O*(*mn*) time. The binary tree with the highest score is then the ML binary genealogical tree which directly provides the ML mutation tree with attachments when we change the representation back to mutations trees [Fig. 1(f)].

We can search the binary tree space with analogous moves as for the mutation trees. A *prune and reattach* move can be performed by detaching one half of any internal binary divide (the remaining neighbouring edges join together) and reinserting it into any of the edges then present. We can also *swap leaf labels*. The size of the relevant binary tree space is 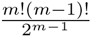 whichmaybesmallerthanthemutation tree space of (*n* + 1)^(*n*–1)^, or easier to search, allowing the ML tree to be discovered more quickly. This alternative representation is implemented in the SCITE software package.

In the binary tree space, one can further marginalise, but this is over the mutation placement rather than the sample attachments and the resulting posterior distribution does not translate directly into one over the mutation tree space.

## Software availability

SCITE has been implemented in C/C‫‫ and is freely available under a GPL3 license at https://github.com/cbg-ethz/SCITE.

## Additional file descriptions

Supplementary figures, the supplementary table and a description of the additional MCMC moves and their effect on the convergence is included in the supplementary pdf file ‘`Supplementary_material.pdf’"`.

## Additional information

This paper was selected for oral presentation at RECOMB 2016 and an abstract is published in the conference proceedings.

## Availability of data and materials

The published datasets analysed are available from the supplementary material of Hou et al. (2012); Xu et al. (2012) and in Figure 2f of Wang et al. (2014). The data matrices are also included with SCITE.

## Ethics

Ethics approval is not applicable for this study.

## Author’s contributions

KJ and JK designed and implemented the method. KJ, JK and NB wrote the manuscript.

